# Mapping neural subspace dynamics onto the structure of the mouse descending motor system

**DOI:** 10.1101/2025.10.17.682917

**Authors:** Munib A. Hasnain, Yujin Han, Michael N. Economo

**Affiliations:** Department of Biomedical Engineering, Boston University, Boston, M; Center for Systems Neuroscience, Boston University, Boston, MA; Center for Neurophotonics, Boston University, Boston, MA

## Abstract

The motor cortex supports various cognitive and motor functions. To prevent interference between these processes, the associated neural dynamics may be organized into orthogonal subspaces. In this ‘subspace model’, activity in a ‘movement-null’ subspace encodes internal processes, while activity in a ‘movement-potent’ subspace relates to movements. The biological implementation of this model – how activity in different subspaces map onto neural circuits – remains unclear. Particularly, it is unknown whether different cell types, with specific connectivity patterns, preferentially contribute to specific subspaces. Here, we test whether the cell type that directly links motor cortex to motor centers in the medulla and spinal cord – lower layer 5b extratelencephalic neurons (L-ETN) – preferentially encodes activity contained in the movement-potent subspace. We performed cell-type-specific recordings in the motor cortex while mice performed a delayed-response licking task, and decomposed population activity into movement-null and movement-potent subspaces. We find that L-ETN activity spans both movement-null and movement-potent subspaces in a manner that could not be distinguished from the broader motor cortex population. Notably, downstream medullary circuits retain a subset of the movement-null dynamics contained in motor cortex, with select movement-null signals specifically filtered out. These results indicate that a distinct cortical output alone does not mediate the cancelation of ‘null’ dynamics; movement-null computations are instead distributed across the descending motor hierarchy.

## Introduction

In mammals, the motor cortex (MCtx) is responsible for a broad range of processes that are essential for movements. For example, in both primates and rodents, the motor cortex is essential for planning upcoming movements^1–6^, learning to produce new movements^7–9^, associating movements with sensory cues^10–12^, and maintaining their accuracy via feedback control^13–17^. How these diverse processes are carried out simultaneously within the circuitry of the MCtx has been the subject of much experimental and theoretical interest^18–21,21^.

Individual neurons often contain activity patterns related to multiple processes^5,22–24^. For example, single MCtx neurons may encode sensory stimuli^25,26^, upcoming actions^22,27^, kinematic parameters of movement^28,29^, reward signals^30,31^ – or some mixture thereof^5,22,32^ – suggesting that separate processes are not mediated by separate neuronal subpopulations.

An alternative explanation posits that the neural dynamics supporting separate processes are not segregated in different neural populations, but in orthogonal subspaces of neural activity^19,21,33,34^. In this framework, population activity at each moment in time is taken to be a point in a high-dimensional state space where each axis represents the firing rate of one neuron^6,35^. A ‘subspace’ is spanned by any smaller set of directions in state space – each direction a weighted average of neural activity determined, for example, by dimensionality reduction^36,37^. When two subspaces are orthogonal to one another, the portion of neural activity contained within one subspace may evolve in a manner that is independent of its evolution in the other, allowing a circuit to subserve different functions without interference. This ‘subspace model’ also suggests a mechanism for selectively controlling the flow of information between connected brain regions^21,38^. Specific linear combinations of population activity, together forming an ‘output-potent’ subspace may shape neural dynamics in a connected brain region^21,38^. Other combinations of neural activity, defining an ‘output-null’ subspace, do not influence activity in that connected region, but instead may contain dynamics that support local computations^21,34^. By extension, specific linear combinations of neural activity may relate directly to the execution of movements, forming a ‘movement-potent’ subspace^19,21,39^. The orthogonal complement of the movement-potent subspace is a ‘movement-null’ subspace, which contains dynamics not directly related to ongoing movements^19,21,39^.

Several experimental observations support the subspace model. Dynamics related to motor planning and movement execution appear confined to orthogonal subspaces^19,21^, rather than separate groups of neurons. In monkey M1, posture-related and goal-related components of population activity occupy orthogonal subspaces, allowing for flexibility in the integration of posture across many behaviors^40^. Learning-related changes in premotor cortex evolve within an output-null subspace, along dimensions orthogonal to those that influence M1 activity, allowing preparatory activity to be modulated without altering dynamics that affect downstream circuits^41^. Further, artificial neural networks trained to solve common behavioral tasks spontaneously learn to segregate separate signals into orthogonal subspaces, rather than segregated subnetworks^19,42,43^.

In the subspace model, all neurons are treated equally analytically – every neuron can contribute to every subspace and every neuron is equally capable of influencing dynamics in all connected regions^19,21^. However, like many neural circuits, the anatomy of the MCtx is highly structured^44–47^. Cell types are stratified by cortical layer^44,45^ and have characteristic local and long-range connectivity patterns^45,47,48^. Despite the expanding scope and influence of the subspace model^33^, it remains unclear if and how this computational abstraction maps onto real neural circuits^5,49^. Do different cell types contribute to specific subspaces? Where are projections of activity into different subspaces computed? Until these and other questions are answered, the biological significance of the subspace model will remain unclear.

In the motor cortex, extratelencephalic neurons (ETN; classically termed ‘pyramidal tract neurons’) project to a diverse set of subcortical motor centers^46,48,50–52^. In mice, ETNs in the lower half of layer 5b (L-ETN) represent a genetically distinct cell type^53^ (Allen Brain Cell Atlas cell type ‘0364 L5 ET CTX Glut_2’^44^) that provides the only direct cortical connection to the motor circuits in the medulla and spinal cord that drive movements^48,53^. L-ETNs are thus anatomically positioned to encode and communicate cortical motor commands – contained within the movement-potent subspace – to downstream motor circuits as the last stage of cortical processing. Although previous work in rodents and primates has suggested that ETNs also contain output-null preparatory dynamics in addition to motor commands^1,53–56^, these studies have been unable to differentiate L-ETNs from ETNs with other functions^53^, have not tracked animals’ movements comprehensively enough to accurately identify preparatory dynamics^39,57^, or both (see ***Discussion***).

Here, leveraging cell-type specific electrophysiology and an analytical procedure we developed to accurately estimate movement-related subspaces in mice^39^, we test the hypothesis that L-ETNs preferentially express dynamics contained within the movement-potent subspace. This hypothesis would imply that L-ETNs compute projections of MCtx dynamics onto movement-potent dimensions to produce motor commands. Surprisingly, we find that the dynamics of L-ETNs span both movement-null and movement-potent subspaces in a manner that could not be distinguished from other neurons in the MCtx. We further find that motor circuits in the medulla, directly postsynaptic to L-ETNs, contain only a subset of the output-null dynamics encoded by L-ETNs, indicating the site at which a critical computation – the ‘canceling out’ of null dynamics along the descending motor pathway – is carried out.

## Results

### Cell-type specific recordings of MCtx dynamics during movement

Water-restricted mice were trained to perform a delayed-response licking task. One of two auditory tones (3 kHz or 12 kHz) was presented during a sample epoch indicating reward location (left or right). Following a brief delay epoch, a go cue instructed mice to perform a directional tongue movement^10,58,59^ (**Fig. 1a-c**; *n* = 9 mice, *n* = 27 sessions). During behavior, we performed extracellular recordings from the tongue/jaw primary motor cortex (tjM1) and anterolateral motor cortex (ALM; putative murine secondary motor cortex) in expert mice (**Fig. 1d**, *bottom*). Both cortical regions exhibit diverse activity patterns related to motor planning and movement execution^10,11,60^. L-ETNs in both regions (ALM: *n =* 49; tjM1: *n =* 55) were identified through photostimulation of their axon collaterals in the reticular nuclei of the medulla, which directly mediate orofacial movements^53,61–63^ (**Fig. 1d**, *top*). Direct axonal projections to the medullary reticular nuclei, a defining characteristic of L-ETNs in the ALM and tjM1, were confirmed through spike collision tests^64^ (**Fig. 1g**). Other recorded units were labeled as unidentified neurons (UNs; ALM: *n =* 1020; tjM1: *n =* 830).

**Figure 1:**
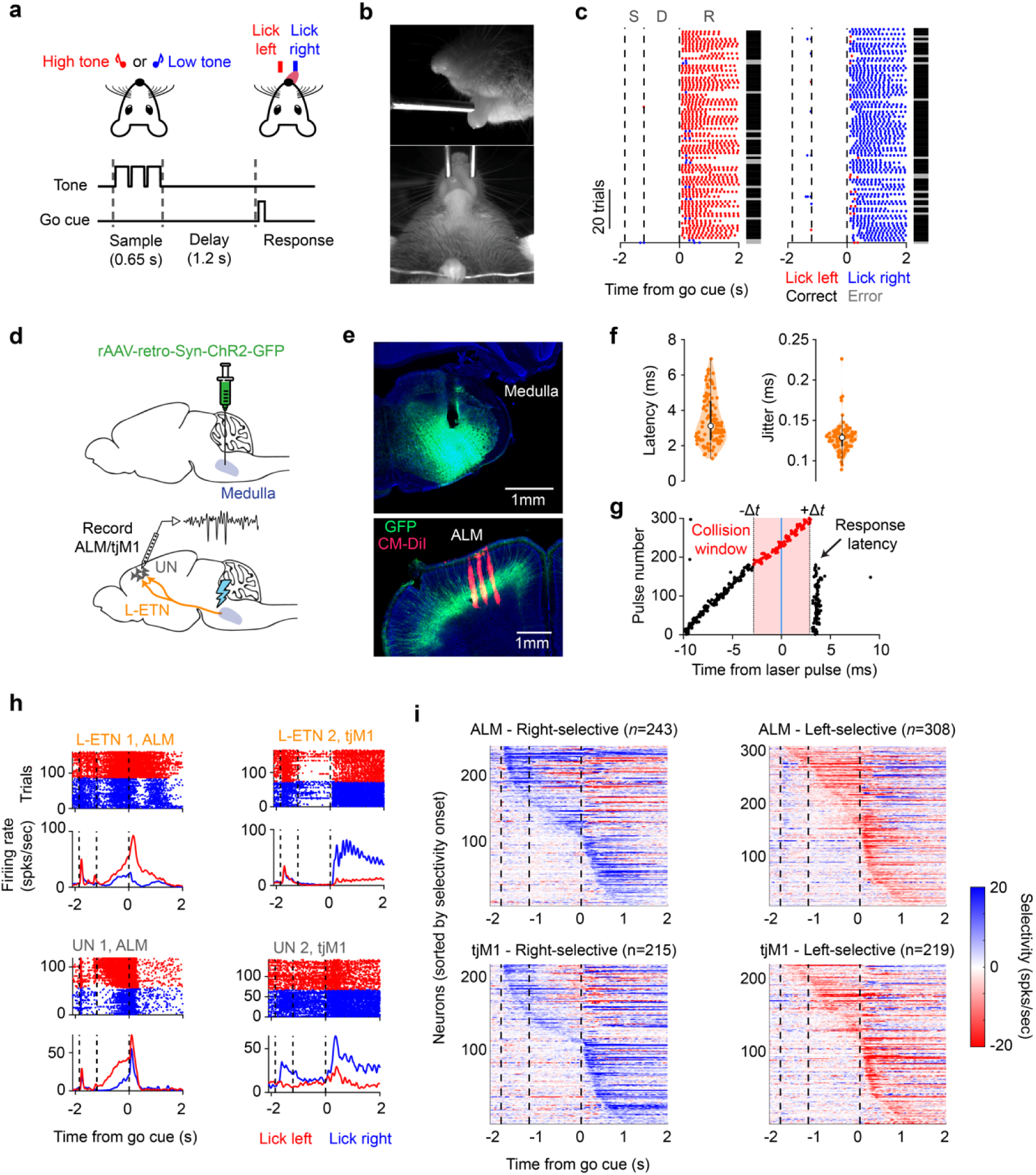
Electrophysiological recordings of L-ETNs during behavior. **a**, Schematic of the delayed-response licking task. An auditory stimulus during the sample epoch (0.65 s) instructed mice to lick for reward to the right (2 kHz tone; random 50% of trials) or left (8 kHz; 50%). Mice were trained to withhold licking during a delay epoch (1.2 s) and initiate a response following an auditory go cue. **b**, High-speed (400 Hz) videography from side and bottom views. **c**, Performance in an example session, separated by trial type. Dots indicate lickport contacts to the right and left. Black and gray bars indicate correct and error trials, respectively. Dashed lines indicate the onset of the sample epoch, delay epoch, and go cue. S: sample, D delay, R: response. **d**, Schematic of L-ETN targeting (*top and bottom*) and silicon probe recording (*bottom*) strategies. **e**, Expression of ChR2 (GFP) and optic fiber placement in the medulla (*top*). Expression of ChR2 in L-ETNs in the ALM and dye-labeled silicon probe tracks (CM-DiI) (*bottom*). Example histological sections from the same mouse are shown. **f**, Latency and jitter (s.d. of latency) of L-ETN responses following the onset of axonal photostimulation (*n* = 104 L-ETNs, *n* = 9 mice, *n* = 27 sessions). Mean and interquartile range indicated. **g**, Spike collision test for an example L-ETN. Trials were binned (0.5 ms) according to the timing of spontaneously occurring spikes. Photostimulation-evoked spikes are absent at the expected latency (+Δ*t*) when a spike occurred spontaneously within the collision window (−Δ*t* to +Δ*t*). **h**, Spike rastergrams (*above*) and trial-averaged firing rates (*below*) for two L-ETNs and two UNs. **i**, Selectivity of units in the ALM (*top row*) and tjM1 (*bottom row*). Units that were selective (*p* < 0.05; shuffle test) during at least one task epoch are shown.

L-ETNs also exhibited diverse activity patterns, in many cases selective for a particular trial type (left or right) during one or more task epochs (**Fig. 1h,i**;). The proportion of L-ETNs that exhibit trial-type selectivity in each of the task epochs was similar to that of the UN population (sample: 24% of L-ETNs, 20% of UNs; delay: 31% of L-ETNs, 34% of UNs; response: 60% of L-ETNs, 51% of UNs). This parity may indicate that L-ETNs are engaged across preparatory and motor phases of this task. However, mice commonly express choice-specific uninstructed movements (often eye, whisker, nose, and other postural and facial movements) across task epochs^39,57,65^. Preparatory dynamics predictive of upcoming instructed movements (lick left/right) can be difficult to differentiate from dynamics related to the execution of ongoing choice-specific uninstructed movements^39,65^. However, these dynamics should evolve in separate subspaces, according to the subspace model, with preparatory dynamics contained within the movement-null subspace, and dynamics related to ongoing uninstructed movements contained within the movement-potent subspace^39^.

### Subspace specificity of L-ETNs

To test whether L-ETN activity was contained principally within the movement-potent subspace, we decomposed single-trial neural activity into orthogonal movement-null and movement-potent subspaces in a manner that accounts for both instructed and uninstructed movements^39^ (**Fig. 2a,b**). Briefly, movement-null and movement-potent subspaces were jointly identified as the linear combinations of neural activity that maximally captured neural variance during periods of animal stationarity and movement, respectively (see ***Methods***). Defined in this way, neural activity related to internal processes, such as motor planning and reward processing should be contained within the movement-null subspace, while activity related to ongoing movements, including sensory reafference, proprioceptive signals, and motor commands, comprise the movement-potent subspace^39^. This analytical approach permits dynamics related to movements and internal processes to be expressed during any task epoch, and to co-occur simultaneously^39^. We found that the movement-null and movement-potent subspaces contained similar proportions of neural variance and together captured 78% of the variance in single-trial neural activity (**Fig. 2c**; movement-null: 37 ± 7.5%, mean ± s.d.; movement-potent: 41 ± 6.9%; *n* = 27 sessions).

**Figure 2.**
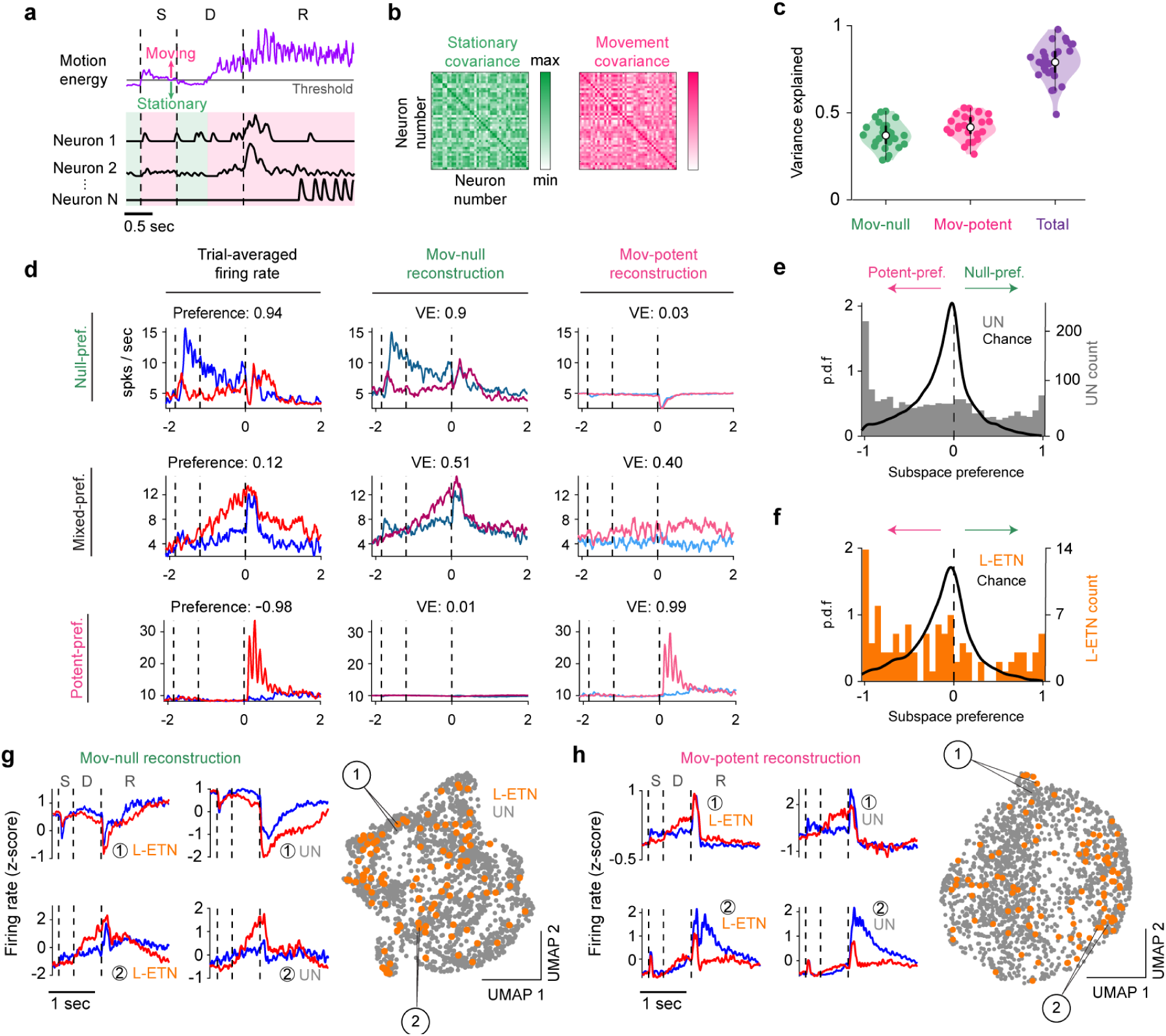
– Neural activity within movement-null and movement-potent subspaces. **a**, Schematic for estimation of moving (*magenta*) and stationary (*green*) time points on single trials. S: sample, D delay, R: response. **b**, Covariance matrices of single-trial neural activity from stationary (*left*) and moving (*right*) time points for an example session used to estimate movement-null and movement-potent subspaces. **c**, Variance of single-trial neural activity explained by the movement-null (*green*), movement-potent (*magenta*), and both subspaces (*purple*). Points indicate sessions (*n* = 27). Mean and interquartile range indicated. **d**, Trial-averaged firing rates (*left column*) for three example units (*rows*). Trial-averaged rates reconstructed from projections of activity into the movement-null (*middle*)and movement-potent subspaces (*right*). *Preference*: subspace preference index, *VE*: variance explained. **e**, Subspace preference index distributions for UNs. **f**, Same as (**e**) for L-ETNs. **g**, UMAP plot of neural activity contained within the movement-null subspace (*right*). The movement-null activity of adjacent units is shown to highlight their similarity. **h**, Same as (**f**), but for activity within the movement-potent subspace.

**Extended Data Figure 1.**
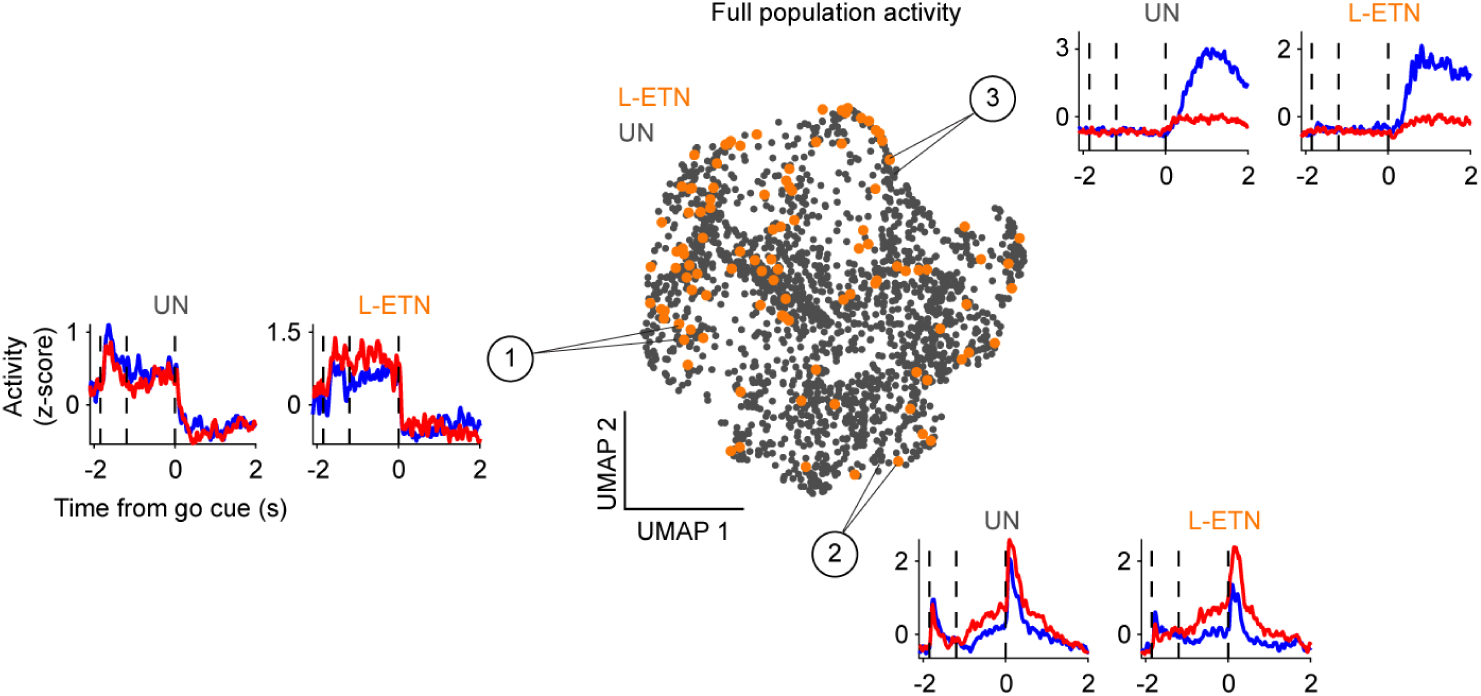
– UMAP applied to the full population activity. UMAP projection of trial-averaged neural activity prior to subspace decomposition. The z-scored firing rates of three pairs of adjacent L-ETNs and UNs are shown.

Individual units can exhibit dynamics contained within the movement-null subspace, the movement-potent subspace, or both. We quantified the relative amounts of each unit’s activity within the two subspaces by computing a subspace preference index, defined as the difference in variance captured by the two subspaces normalized by the sum of the variance captured by both subspaces. A subspace preference of +1 (−1) indicates that a unit’s variance is primarily captured by the movement-null (movement-potent) subspace (**Fig. 2d**; *top, bottom*). A near-zero subspace preference indicates that both subspaces capture similar amounts of variance (**Fig. 2d**, *middle*). As in previous work, we observed an unusually large number of units with activity contained almost entirely within the movement-potent subspace (**Fig. 2e**), indicative of a population MCtx neurons with activity related predominantly to the execution of movements^39^. Because L-ETNs represent the direct conduit for cortical motor commands to brainstem and spinal motor nuclei, we tested the hypothesis that their dynamics may be principally contained within the movement-potent subspace. This hypothesis implies that linear combinations of MCtx activity responsible for driving movements are computed in L-ETNs. Surprisingly, we found that the subspace preferences of L-ETNs could not be distinguished from UNs (**Fig. 2f**; *p* = 0.30, two-sample *t*-test).

While subspace preferences reveal the relative proportions of movement-null and movement-potent dynamics in cells of each population, they do not capture the structure of their underlying activity patterns. We next asked whether L-ETN activity spans the collection of task-related activity patterns observed within each subspace. We reconstructed L-ETN and UN trial-averaged neural activity from the movement-null and movement-potent subspaces, independently, and projected these reconstructions into a two-dimensional space using UMAP (**Fig. 2g,h**). Each point in the embedding represents the full temporal profile of a single unit’s activity within a given subspace. L-ETNs populated the full extent of both subspaces, albeit not necessarily uniformly (**Fig. 2g,h**). Thus, while some activity patterns may be enriched in L-ETNs, L-ETN dynamics span the full set of temporal motifs represented within each subspace.

Together, these findings demonstrate that the activity of L-ETNs is not preferentially contained within the movement-potent subspace. Instead, L-ETNs exhibit dynamics spanning the entirety of both the movement-null and movement-potent subspaces, in a manner that was not qualitatively different than other cortical neurons. This suggests that L-ETNs propagate a broad set of information to downstream motor circuits, including signals not directly related to motor output.

### Similarity in subspace preference is robust to false-negatives

The proportion of L-ETNs identified in our recordings (104/1954, 5.3%) is consistent with the overall prevalence of L-ETNs in the motor cortex (2.5-7%) ^44^, although we sampled cortical layer 5 more densely than other layers (**Extended Data** Fig. 2b). Therefore, it is possible that some L-ETNs were not identified via phototagging (e.g. due to lack of opsin expression) and were therefore classified as UNs. Misclassifying L-ETNs as UNs could cause the subspace preferences for the two populations to appear more similar than if L-ETNs were fully excluded from the UN population.

To identify potential false negatives, we used waveform shape and firing rate statistics to discriminate UNs from L-ETNs using linear discriminant analysis (LDA; **Extended Data** Fig. 2c,d). Each unit’s feature vector was projected onto the discriminant boundary, and we defined an “L-ETN-like” criterion as UNs that exceeded the 5^th^ percentile of the L-ETN distribution (**Extended Data** Fig. 2e). We further restricted “L-ETN-like” units to those in or near lower layer 5b (see ***Methods***). Applying these criteria yielded 140 L-ETN-like UNs. Excluding these from the UN population did not qualitatively alter any results. The subspace preferences of L-ETNs remained indistinguishable from UNs (**Extended Data** Fig. 2f; *p* = 0.22, two-sample *t*-test).

**Extended Data Figure 2.**
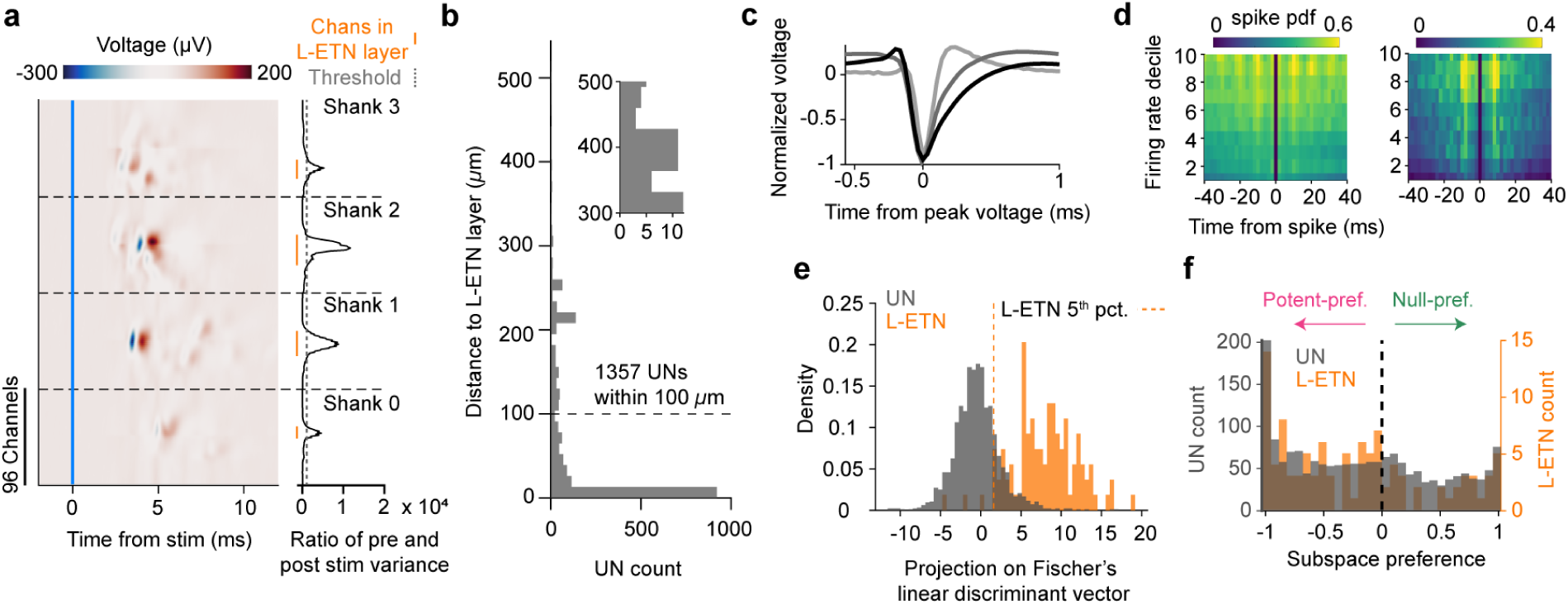
– Controlling for false negatives L-ETNs included in UN population. **a**, Mean extracellular voltage in response to photostimulation across sites in an example recording using a four-shank Neuropixels 2.0 probe (*left*). Ratio of the variance in mean voltage following (0 to +10 ms) and preceding (−5 to 0 ms) photostimulation (*right*). **b**, Distance of UNs (*n* = 1850) to the L-ETN layer. *Inset*, expanded view of distribution at distances between 300 and 500 μm. **c**, Average normalized waveform of three example units. **d**, 3D-autocorrelogram (ACG) for two example units. Each row contains a 2D-ACG for a different decile of mean firing rate. **e**, Projection of UN and L-ETN features onto Fischer’s linear discriminant vector. The 5^th^ percentile of the L-ETN distribution, used as a threshold for identifying UNs that were similar to ‘L-ETN,’ is shown. **f**, Subspace preference distributions of L-ETNs (*right axis*) and UNs (*left axis*) after excluding UNs that were both (1) within 100 μm of the L-ETN layer and (2) exhibited electrophysiological features similar to L-ETNs.

### Investigating movement encoding using a nonlinear model

The subspace model explicitly assumes that *linear* combinations of neural activity encode internal variables and effect movements^19,21,33,39^. Alternatively, projections of neural activity onto nonlinear manifolds may better describe the relationship of neural activity to behavior^66,67^. To address the possibility that L-ETN dynamics relate to movements nonlinearly, we trained artificial neural networks to predict single trial neural firing rates from high-speed videography^68^ (**Fig. 3a,b**). Neural network models explained 27.9 ± 20.2% (mean ± s.d., *n* = 1954 neurons) of the variance in MCtx activity, outperforming linear regression models (10.6 ± 17.9%; **Fig. 3c**; *p* = 1.1 × 10^−10^, paired two-sided *t*-test), and comparable to the accuracy of previous neural-network based motor cortex encoding models^58,68,69^.

**Figure 3.**
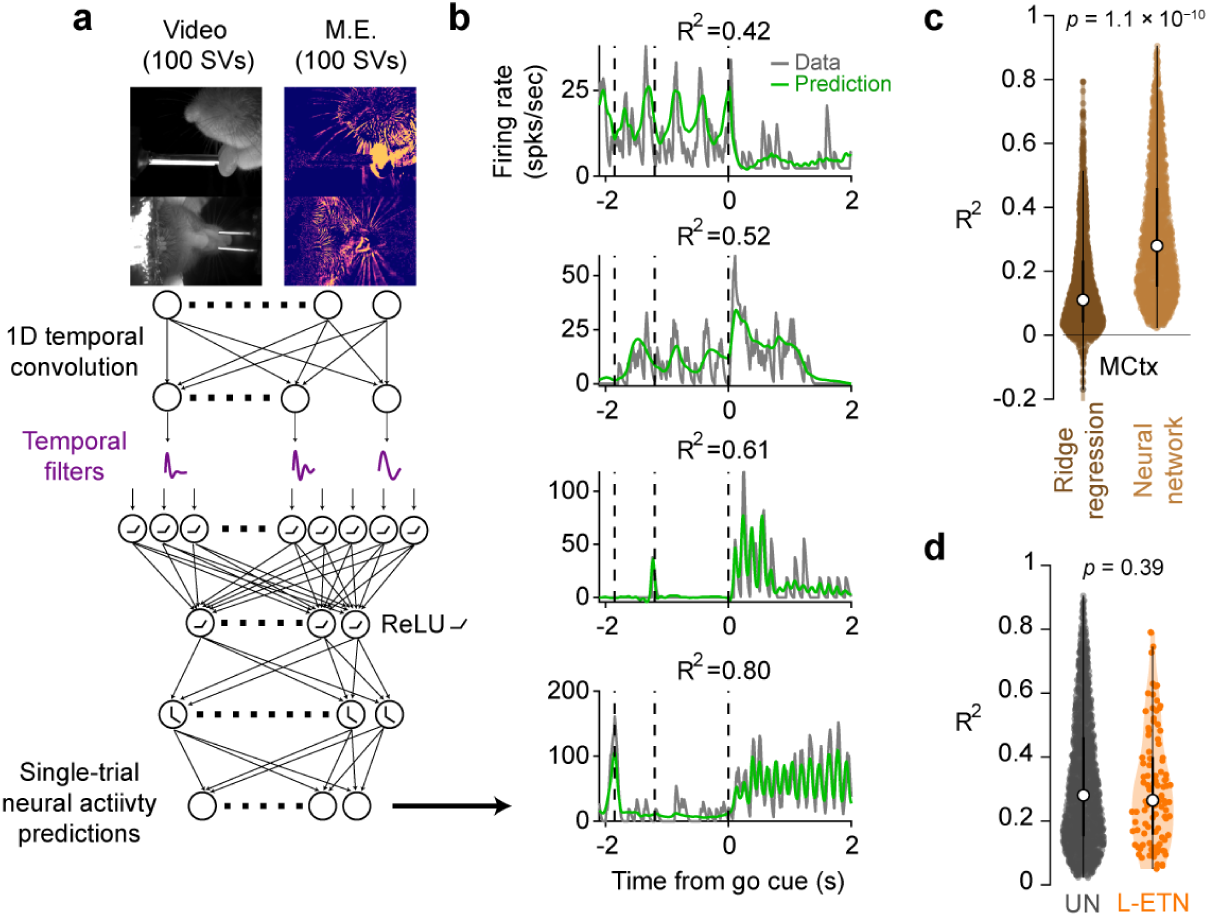
– Single-trial, neural network prediction of neural activity. **a**, Schematic of neural network used to predict single-trial neural activity from singular vectors (SV) of video and motion energy (ME) movies. **b**, Actual and predicted firing rates on example trials for four units. **c**, Variance of MCtx single-trial neural activity explained by ridge regression or neural network model (*n* = 1945 units). Mean and interquartile range indicated. **d**, Variance of L-ETN (*n* = 104) and UN (*n* = 1850) activity explained by neural network model. Mean and interquartile range indicated.

We examined whether the activity of ETNs could be better explained by animals’ movements than UNs using this nonlinear neural network model. Again, L-ETNs could not be distinguished from UNs in terms of how well their dynamics were explained by movements (L-ETN: 26.4 ± 17.9%, mean ± s.d., UN: 28.0 ± 20.1%; *p* = 0.39, two-sample *t*-test). Unexplained variance in the activity of ETNs and UNs may relate to internal processes unrelated to ongoing movement (**Fig. 3d**). This result further supports the conclusion that L-ETNs contain both movement-null and movement-potent dynamics, and that the relative strength of these components is similar to that observed in other cortical neurons.

### Coding specificity of L-ETNs

Our previous work established that ETNs can be divided into two major classes: L-ETNs and U-ETNs, with the latter type located in upper layer 5b^53^. Among these, only L-ETNs were found to express dynamics that preceded movement initiation, consistent with a direct role in driving movements^53^. In contrast, only U-ETNs were found to persistently encode the identity of animals’ upcoming movements, suggesting a role in motor planning^53^. This cell-type specialization seemingly contrasts with the observation that L-ETNs broadly encode and communicate both movement-null and movement-potent signals to downstream regions. Here, we find similar features in the activity of L-ETNs (**Extended Data** Fig. 3). L-ETN dynamics changed rapidly at the go cue, just preceding movement initiation, whereas the UN state switch was slower and followed movement onset (**Extended Data** Fig. 3b-d; *p* = 7.6 × 10^−18^, paired *t*-test between slope of curves in **Extended Data** Fig. 3d; movement onset: 105 ± 36 ms following the go cue, mean ± s.d.; *n* = 27 sessions). Further, we found that the identity of upcoming movements was stably encoded in UNs, but not L-ETNs (**Extended Data** Fig. 3a). Thus, while the dynamics of L-ETNs (1) span movement-null and movement-potent subspaces in a similar manner to UNs, and (2) span the set of dynamics contained within each subspace, finer-grained analysis reveals specific features of their activation patterns that are not shared by the larger UN population.

**Extended Data Figure 3.**
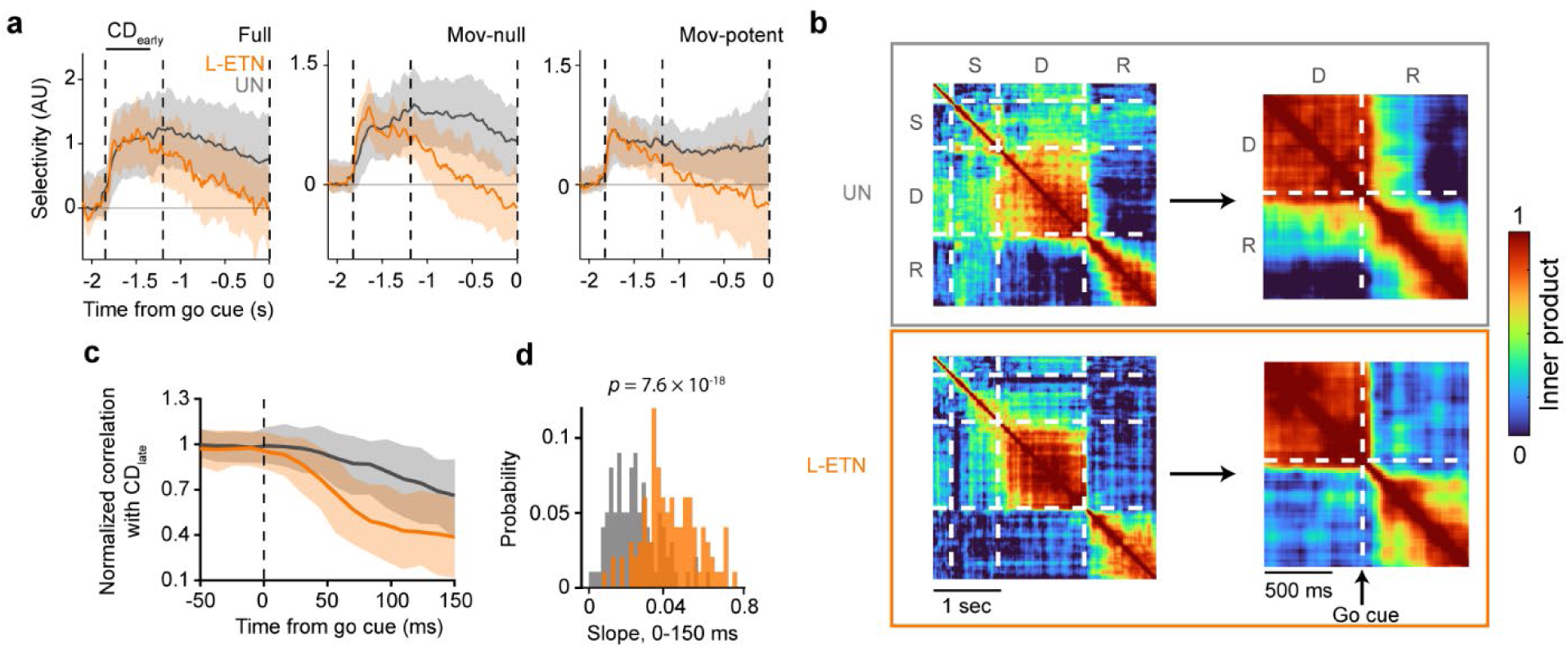
– Coding specificity of L-ETNs. **a**, Time course of selectivity along the coding direction (CD) that best encodes upcoming movements following stimulus onset, calculated from the full space of neural activity (*left*) and activity contained in the movement-null (*middle*), or movement-potent subspace (*right*). The CD was defined as the vector in neural activity state space that best separates lick left and lick right trajectories in the first 500 ms of the sample epoch. While projections of UN activity along the CD were selective until the go cue, the selectivity of L-ETN activity along the CD decayed to baseline following the sample epoch, recapitulating previous observations. Shaded regions denote 5-95% confidence intervals (denoting region significantly greater than zero, *p* < 0.05 one-sided test, bootstrap, *n* = 1,000 iterations). **b**, Heatmaps of the similarity (inner product) between CDs calculated at different time points. **c**, Correlation between the CD determined at each time point and the CD observed prior to (−500 ms to 0 ms) the go cue. A rapid change in the CD occurs following the go cue in L-ETNs, starting approximately at the time of movement initiation, recapitulating findings in prior work. This switch in dynamics was not observed in UNs. Shaded areas represent the 95% confidence interval of the bootstrapped distribution (*n* = 1,000 iterations). **d**, Distribution of slopes across bootstrap iterations (rate of change of correlation) observed in (**c**) from 0 to 150 ms following the go cue. The slope was larger for L-ETNs than UNs (*p* = 7.6 × 10^−1^^8^, paired *t*-test).

### A subset of null dynamics is found downstream of motor cortex

By definition, movement-null dynamics do not drive movements, and therefore must not be propagated to the motor periphery. Rather, they must be ‘filtered out’ somewhere along the descending pathway so as not to drive changes in the activation of motor neurons. We found that movement-null dynamics are robustly encoded by L-ETNs, and therefore not filtered out at the level of cortical output. We next examined whether movement-null dynamics are communicated to the medullary reticular nuclei, which are directly postsynaptic to L-ETNs and directly presynaptic to orofacial motorneurons^61,62^. We analyzed the publicly available mesoscale activity map (MAP) dataset, in which simultaneous recordings were made from both the ALM and medullary targets of L-ETNs while mice performed the same delayed-response task^58^. In each session, we computed subspaces independently in each region (*n* = 16 sessions, ALM: 604 units, medullary reticular nuclei: 724 units; see ***Methods***).

Individual neurons in the medullary reticular nuclei exhibited activity patterns representative of both movement-null and movement-potent dynamics (**Fig. 4a**). Many units exhibited selectivity for either right or left licks in different task epochs, although selectivity was most prominent during the response epoch (sample: 7%; delay: 21%; response: 49%; *n* = 724 units **Fig. 4b**). In the MAP dataset, the variance explained by movement-null and movement-potent subspaces in the ALM was similar to that reported here, with both subspaces explaining comparable amounts of variance (movement-null: 36.0 ± 7.40%, mean ± s.d.; movement-potent: 37.1 ± 8.70%; *p* = 0.15, paired two-sample *t*-test; *n* = 16 sessions; **Fig. 4c**). In contrast, in the medulla, the movement-null subspace explained less variance than the movement-potent subspace (movement-null: 26.4 ± 3.90%, mean ± s.d.; movement-potent: 44.8 ± 7.12%; *p* = 1.9 × 10^−9^, paired two-sample *t*-test; *n* = 16 sessions; **Fig. 4c**, right). This suggests that specific movement-null signals may be filtered out or that movement-null signals may be attenuated nonspecifically by medullary circuits.

**Figure 4.**
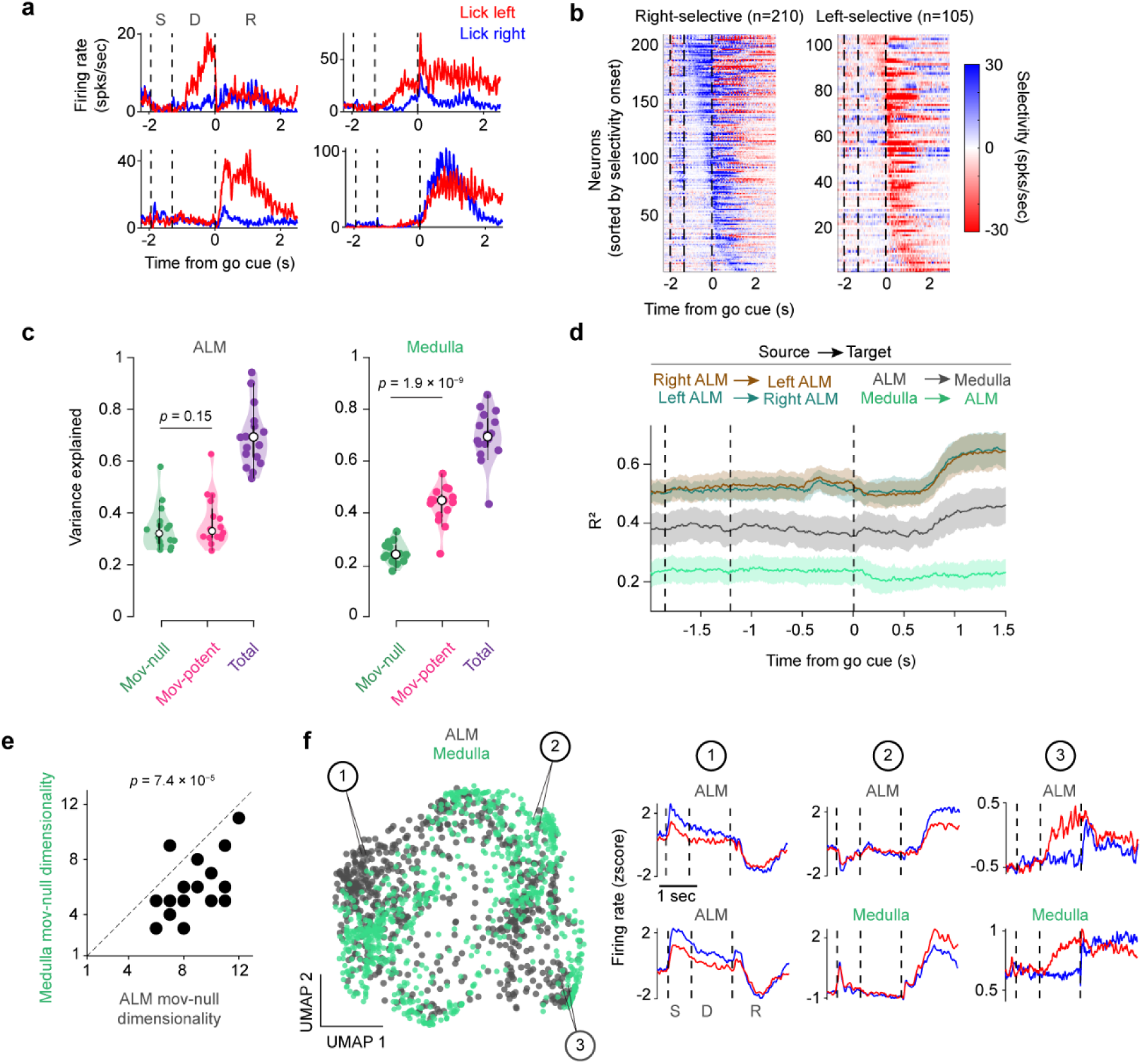
– Movement-null dynamics in the medullary reticular nuclei. **a**, Trial-averaged firing rates of four example units in the medullary reticular nuclei. S: sample, D delay, R: response. **b**, Selectivity of units with significant selectivity during at least one task epoch. **c**, Variance of single-trial neural activity contained within the movement-null (*green*), movement-potent (*magenta*), and both subspaces (*purple*; *n* = 16 sessions; paired two-sample *t*-tests). Mean and interquartile range indicated. **d**, Variance explained (R^2^) for cross-validated predictions of single-trial movement-null activity in target regions from single-trial movement-null projections in source region. Mean and 95% confidence intervals indicated. **e**, Dimensionality of the movement-null subspace in the ALM versus the medulla. **f**, UMAP plot of neural activity contained within the movement-null subspace (*left*). The movement-null reconstructions of two example ALM units (*left column*) are shown to highlight activity patterns only present in the ALM movement-null subspace. Middle and right columns show the activity of units in the ALM and medulla that are adjacent in the UMAP embedding. S: sample, D: delay, R: response.

To determine whether medullary movement-null dynamics could be inherited from ALM, we performed cross-area decoding using regularized linear regression. We predicted movement-null activity in the medullary reticular nuclei at each time point from the previous 40 ms of movement-null activity in the ALM (**Fig. 4d**). For reference, we applied the same analysis to cross-hemisphere ALM recordings (*n* = 10 mice; *n* = 20 sessions; *n* = 1116 neurons). We selected ALM-to-ALM movement-null decoding as an upper bound because the two hemispheres are densely interconnected^4,27^, and perturbations to one hemisphere during working-memory tasks are recovered by information from the other hemisphere^4^. Medullary movement-null activity could be predicted from ALM movement-null dynamics (40 ± 8.1%, mean ± s.e.m; **Fig. 4d**) nearly as accurately as from the contralateral ALM (54 ± 7.0%, mean ± s.e.m; **Fig. 4d**). However, this relationship was asymmetric. ALM movement-null dynamics were less accurately predicted from movement-null dynamics in medullary reticular nuclei (21 ± 6.7%, mean ± s.e.m; *p* = 1.2 × 10^−4^, paired *t*-test comparing ALM-to-medulla and medulla-to-ALM predictions; *n* = 16 sessions; **Fig. 4d**). This asymmetry suggests that a subset of the movement-null dynamics in the ALM are represented in medullary reticular nuclei, despite their robust encoding in L-ETNs.

To further distinguish between a global attenuation of cortical movement-null signals by medullary circuits versus selective filtering of specific signals, we compared the dimensionality of movement-null dynamics in each region. If movement-null signals were nonspecifically attenuated, the distribution of variance across eigenvectors of movement-null dynamics would be preserved, yielding similar dimensionality across regions. If specific movement-null signals were selectively filtered, some eigenvectors should capture disproportionately less (or more) variance, resulting in a smaller effective dimensionality^70^. Consistent with the latter, the dimensionality of the movement-null subspace was lower in medullary reticular nuclei than in ALM (**Fig. 4e**; *p* = 7.4 × 10^−5^, paired *t*-test). Further, examining a UMAP embedding of ALM and medullary movement-null activity, we found that the activity patterns observed in the medullary nuclei did not span the collection of movement-null activity patterns observed in the ALM (**Fig. 4f**), supporting a selective communication of movement-null signals to the medullary nuclei.

Together, these findings suggest that movement-null dynamics are distributed along the descending pathway: motor cortical ‘null’ dynamics are indiscriminately communicated to medullary reticular nuclei, where a subset of those signals are specifically filtered. This implies that the remaining movement-null dynamics in medullary reticular nuclei are either not transmitted to orofacial motor neurons, or are canceled by motor neurons that predominantly integrate activity within the movement-potent subspace.

## Discussion

Here we examined two questions related to the biological implementation of the subspace model: (1) do different cell types contribute to specific subspaces of neural activity and (2) where are projections of activity onto those subspaces computed? We addressed these questions by performing cell-type-specific recordings of L-ETNs in mice performing a delayed-response licking task (**Fig. 1**). L-ETNs, a single cell type that forms the only direct projection from the MCtx to medullary and spinal motor centers, are anatomically positioned to communicate motor commands to downstream motor circuits and therefore represent the most likely candidate cell type to specifically encode movement-potent dynamics. We decomposed single-trial neural activity into a movement-null subspace, containing dynamics related to processes uncoupled from movement, and a movement-potent subspace encoding dynamics related to ongoing motor actions. This decomposition enabled us to quantify the degree to which the activity of individual neurons is contained within each subspace.

A subpopulation of MCtx neurons, substantially larger than expected by chance, had activity almost exclusively captured by the movement-potent subspace (**Fig. 2e,f**), lending plausibility to the idea that the dynamics of one or more cell type(s) may evolve preferentially within a particular subspace. Surprisingly, however, we found that the activity of L-ETNs was not only not specifically contained within the movement-potent subspace but was not even preferentially contained within the movement-potent subspace compared to all other neurons in the MCtx. (**Fig. 2f**). Further, L-ETNs contained dynamical motifs spanning the full space of dynamics comprising both the movement-null and movement-potent subspaces (**Fig. 2f**). At a finer-grained level, however, the dynamics of L-ETNs did exhibit unique features – such as fast onset kinetics accompanying movement initiation – that were not shared by other UNs (**Extended Data** Fig. 3).

In primates and mice, it has previously been demonstrated that ETNs exhibit activity in the absence of instructed movements, during motor preparation^1,53^ and observation^54,55^. These results suggest that ETN dynamics span movement-null and movement-potent subspaces. However, mammals contain multiple classes of ETNs. In the mouse, ETNs can be divided into two types that are distinct anatomically, molecularly, and functionally – forming parallel communication channels through which information is broadcast from the cortex to subcortical motor centers. In previous studies, ETNs have typically been identified on the basis of axonal projections to the pons or medullary pyramids (thus, the common name ‘pyramidal tract’ neurons). Both classes of ETNs exhibit these projections, but only L-ETNs project to motor centers in the medulla and spinal cord. In contrast, ETNs in upper L5b (U-ETNs) target a different set of subcortical structures^45,53,71^, including the thalamus, which is known to have an important role in motor planning. Most previous studies examining preparatory dynamics in ETNs have not differentiated between ETN classes – particularly those conducted in primates, where the number and projection patterns of ETN subtypes remain poorly understood.

Further, many previous studies have not accounted for uninstructed movements when examining MCtx dynamics^57^. Uninstructed movements – changes in posture, gaze, facial expressions, and other ‘fidgets’ that are not explicitly related to completion of a task – are often subtle, highly predictive of neural activity, and, critically, correlated with task variables^39,57,65,72^. Uninstructed movements are common in rodents and primates and account for a large proportion of neural activity previously attributed to internal processes^39,65,73^. For example, delay-period preparatory activity, which precedes and predicts movement identity, can be easily conflated with delay-period activity related to uninstructed movements, which are themselves often predictive of upcoming instructed movements^39,72,74,75^. Although primate studies often ensure a lack of delay period movement with EMG, this is typically limited to a single effector that animals are trained to keep stationary^1,21,54,55,76^. It is challenging for animals to withhold all movements. Even in humans, this requires a high degree of cognitive effort, reducing task performance^77^. ‘Regressing out’ uninstructed movements is equally challenging, as complex postural and facial movements are difficult to parameterize and may relate to neural activity nonlinearly^78^. Decomposing activity into complementary movement related and unrelated subspaces avoids these challenges^19,39^.

In simultaneous recordings in the ALM and medullary reticular nuclei, we observed movement-null dynamics in medullary populations, although the movement-null subspace captured less variance than the movement-potent subspace (**Fig. 4c**). This contrasts with activity in the ALM where the two subspaces explained comparable variance (**Fig. 4c**). Cross-area decoding sharpened this picture: medullary movement-null activity was reliably predicted from movement-null activity in the ALM. In contrast, movement-null dynamics in the medullary nuclei were substantially less predictive of movement-null dynamics in the ALM (**Fig. 4d**). This suggests that movement-null dynamics in the cortex are only partially communicated to motor circuits in medullary nuclei, despite their robust encoding in the L-ETNs that connect these regions. Further analyses revealed that the ALM movement-null dynamics were not indiscriminately attenuated in the medullary nuclei, but that specific movement-null signals were selectively ‘filtered out,’ reducing the dimensionality of the medullary movement-null subspace (**Fig. 4e-f**).

Previous studies in mice, cats, and primates have observed movement-null dynamics in medullary and spinal circuits^69,79–81^. In primates, spinal interneurons display preparatory activity that overlaps in time with preparatory activity in cortical neurons^79,80^, yet most are negatively modulated and only weakly selective for upcoming movement direction compared to preparatory activity in MCtx^82^. Our results revealed that neurons in medullary reticular nuclei encode movement-null dynamics that span a broad subset of the collection of cortical movement-null activity patterns and therefore are not only related to motor preparation (**Fig. 4e-f**). The output neurons of the reticular nuclei, or the motor neurons themselves, must then be computing the projections necessary to cancel the remaining movement-null dynamics prior to them reaching the muscles.

Collectively, our results argue that the computations central to the subspace model are distributed along the descending pathway. Rather than being canceled at the level of cortical output, movement-null dynamics in the MCtx are broadcast via L-ETNs to medullary reticular nuclei. A subset of these movement-null dynamics are present in these nuclei, while others are filtered out. Movement-null activity patterns may endow medullary circuits with the flexibility to perform computations necessary for effective motor control but not explicitly tied to the generation of movements^33,83^. The role of cortical ‘null’ dynamics in downstream regions may be an intriguing focus of further work.

## Methods

### Animals

The study used data collected from 9 mice; 3 male and 6 female animals between 8 and 11 weeks were used. All animals used were either C57BL/6J (JAX 000664) or VGAT-ChR2-EYFP (−/−) (bred in-house; breeders: JAX 014548). Mice were housed in a 12-hour reverse dark/light cycle room (ambient temperature of 68-79° F (20-26° C); humidity of 30-70%) with free access to food in the home cage. Access to water was restricted during behavioral and electrophysiology experiments. Sample sizes were not determined using any statistical tests.

### Surgical procedures

All surgical procedures were performed in accordance with protocols approved by the Boston University Institutional Animal Care and Use Committee. For post-operative analgesia, mice were given ketoprofen (0.1 mL of 1 mg/mL solution) and buprenorphine (0.06 mL of 0.03 mg/mL solution) prior to the start of all surgical procedures. Mice were anesthetized using 4-5% isoflurane and maintained with 1-2% isoflurane. Mice were placed on a heating pad in a stereotaxic apparatus. Artificial tears (Akorn Sodium Chloride 5% Opthalmic Ointment) were applied to their eyes and a local anesthetic (Bupivacaine; 0.1 mL of 5 mg/mL solution) was injected under the skin above the skull. The skin overlying the skull was removed to expose the ALM (AP: +2.5 mm, ML: +1.5 mm), tjM1 (AP: +1.5, ML: +2.5 mm), bregma, and lambda. The periosteum was removed, and the skin was secured to the skull with cyanoacrylate (Krazy Glue) around the margins.

For virus injections, the skull was first thinned in a ∼0.5 mm diameter spot above the IRN (AP – 6.65 mm, ML +1.0 mm). Virus was injected at a depth of 4.4 mm using a manual displacement injector (MMO-220A, Narishige) connected to a glass pipette. The glass pipette (5-000-2005, Drummond Scientific) was pulled to a 30 μm tip (P-2000, Sutter Instruments) and the bevel was sharpened on the exposed disk of a hard drive. Pipettes were loaded with mineral oil and then front-loaded with virus before injection. Pipettes were lowered through the thinned skull to a depth of 4.4 mm and virus was injected at a rate of 10 nL/min. Following injection, the pipette was allowed to sit for 10 minutes before retraction. Following retraction, a fiber optic cannula (MFC_200/245-0.37_5mm_ZF1.25(G)_FLT; Doric Lenses) was implanted 300 μm above the injection depth. Dental cement (Jet™). was used to secure the optic fiber.

A headbar was implanted posterior to bregma and secured with superglue and dental cement. Wells to hold cortex buffer (NaCl 125mM, KCl 5mM, Glucose 10mM, HEPES 10mM, CaCl2 2mM, MgSO4 2mM, pH 7.4) during electrophysiology recordings were sculpted using dental cement and a thin layer of superglue was applied over any exposed bone.

### Mouse behavior

Following surgery, mice were allowed to recover for ∼1 week and then placed on water restriction, receiving 1.0 mL of water per day. Behavioral training started 3 days after beginning water restriction. If animals received less than 1.0 mL of water during training, they were supplemented with additional water in their home cage.

Behavioral training was conducted using custom scripts writing for Bpod r0.5 (https://www.sanworks.io/shop/products.php?productFamily=bpod).

Mice were trained on a delayed-response auditory discrimination task until they reached at least 70% accuracy. At the beginning of each trial, one of two auditory tones lasting 0.65 seconds were played; the tone indicating a ‘right’ trial was a low frequency (3 kHz pulses) pure tone, while the tone indicating a ‘left’ trial was a high frequency (12 kHz pulses) pure tone^58^. Each tone was played three times for 0.15 seconds with 0.2 second inter-tone intervals^58^. The delay epoch (1.2 seconds) started after the completion of the sample tone^58^. Following the delay epoch, an auditory go cue was played (6 kHz pure tone, 100 ms) after which animals were rewarded with ∼3 µL of water for contacting the correct lickport. Inter-trial intervals were randomly drawn from an exponential distribution with a mean equal to 1.5 s. Lickport contacts before the response epoch resulted in the current task epoch restarting. If the animal did not respond within 1 s of the go cue, this was considered an ‘ignore’ trial, although responses were typically registered within 300 ms of the go cue.

### Videography and behavioral analysis

High-speed video was captured (400 Hz frame rate) from two cameras (FLIR, Blackfly® monochrome camera, Model number: BFS-U3-16S2M-CS). One provided a side view of the mouse and the other provided a bottom view.

For each pixel in each frame, motion energy was computed as the absolute difference between the median pixel value over the subsequent 5 frames (12.5 ms) and the median pixel value over the preceding 5 frames (12.5 ms). A single motion energy value for the frame was then obtained by taking the 99th percentile (∼700 pixels) of the pixel-wise motion energy values. Using this approach, motion energy was elevated during pronounced, localized movements, such as whisking, while remaining comparatively low during subtle, widespread motions like passive breathing. For each session, a movement threshold was manually determined. Motion energy histograms within a session consistently exhibited a bimodal distribution, with a prominent, low-variance peak at small motion energy values and a secondary, broader peak at higher values. The threshold was chosen as the value separating these two modes. This procedure reliably detected both brief and sustained movements. Alternative strategies, such as defining a fixed “no movement” baseline for comparison, were less effective due to variability in movement timing across trials.

Keypoint tracking of the tip of the tongue from the side view camera was performed using DeepLabCut^84^ and movement onset times were calculated as the first time the tongue was visible following the go cue.

### Electrophysiology recordings

Extracellular recordings were performed in the ALM and tjM1 (together referred to as MCtx here) using one of two types of silicon probes: Neuropixels 1.0^85^ (*n* = 4 sessions) which have one shank and allow recording from 384 channels arranged in a checkerboard pattern or Neuropixels 2.0 (*n* = 23 sessions) which have four shanks and allow recording from 384 channels. For Neuropixels 2.0^86^ recordings, the first 96 channels on each of the four shanks were used for recordings with a footprint spanning 720 μm by 750 μm. ALM and tjM1 recordings were recorded in the same mice, but on different sessions (*n* = 9 mice, *n* = 27 sessions).

For recordings in MCtx, at least 6 hours before recording, a craniotomy (∼1.5-2.0 mm diameter) was made over ALM (AP: +2.5, ML: +1.5 mm) and tjM1 (AP: +1.5, ML: +2.5 mm). 2-5 recordings were performed on a single craniotomy on consecutive days. Prior to insertion, the first ∼100 µm of each shank was coated with CM-DiI (Thermo-Fisher scientific) to determine recording locations post-hoc. After reaching the desired depth, typically between 900 and 1200 µm (MPM System, New Scale Technologies), probes were inserted another 100 µm and slowly retracted 100 µm to minimize probe drift. Brain tissue was allowed to settle for at least 10 minutes before starting recordings. All recordings were made using SpikeGLX (https://github.com/billkarsh/SpikeGLX, <u>phase3a release</u>).

To phototag L-ETNs^53^, we expressed ChR2(H134R)-GFP in all medulla-projecting cells by injecting 50-100 nL rAAV-retro-Syn-ChR2(H134R)-GFP with a titer of 4.65 × 10^13^ into the right or left medulla (AP –6.65 mm, ML +1.0 mm, DV 4.4 mm). This strategy allows us to unambiguously phototag L-ETNs that project to the medulla as they are the only cell type in cortex that does so^48,53^. Immediately prior to and following recording sessions, a separate phototagging protocol was performed to identify putative L-ETNs (cells that respond to photostimuli with short a consistent latency). During each phototagging session, photostimuli were delivered for five minutes at 4 Hz consisting of 0.1-1 ms pulse widths at 90-100 mW.

### Analysis of electrophysiological recordings

Extracellular recordings were bandpass filtered (300-2000 Hz; second order Butterworth filter) and the common mode on ± 12 sites was subtracted from each channel. For ALM and tjM1 recordings, spike events were detected using JRCLUST^87^ (https://github.com/JaneliaSciComp/JRCLUST) and Kilosort4^88^ (https://github.com/MouseLand/Kilosort). Spikes from putative L-ETNs were extracted from the JRCLUST output and sorted manually using a custom MATLAB program. Spikes from units that did not respond to photostimulation were labeled as coming from an unidentified neuron (UN). UN spikes were obtained from the results of Kilosort4, and assigned a quality label using Bombcell^89^ (https://github.com/Julie-Fabre/bombcell). The automated output of Kilsort4 was provided as input to Bombcell^89^, which provided automated curation based on ISI histogram, separation from other units, stationarity across the session, along with other metrics based on waveform and firing rate statistics. UNs were included if they were labeled as ‘single’ or ‘multi’ by Bombcell and contained no more than 20% of the spike times also found in simultaneously recorded L-ETNs. These UNs were finally subject to manual inspection to remove artifacts using Phy (https://github.com/cortex-lab/phy). Recording sessions were included for analysis if they had at least 20 units with firing rates greater than 1 Hz across the entire session. For ALM, our dataset consisted of 49 L-ETNs and 1020 UNs (415 single units and 605 multi units from 13 sessions). For tjM1, our dataset consisted of 55 L-ETNs and 830 UNs (433 single units and 397 multi units from 14 sessions). Spike collision tests were performed for all putative L-ETNs after manual sorting to confirm axonal projections to the medulla^64^.

### Identifying selective neurons and selectivity onset times

Spike counts for each neuron from random and equal samples of correct right and left trials were binned with 5 ms bin width and smoothed using a causal Gaussian kernel (20 ms s.d.). Selectivity (right vs. left) was statistically tested within pre-defined behavioral epochs (time in seconds relative to the go cue; sample: [-1.8 to –1.2]; early delay: [-1.2 to –0.5]; late delay: [-0.5 to 0]; early response: [0 to 0.5]; late response: [0.5 to 1.5]; **Fig. 1i** and **Fig. 4b**). For each epoch and neuron, firing rates were averaged across bins within the epoch and compared between left and right trials. Direction selectivity was quantified using the area under the receiver operating curve (AUROC), a measure of the probability that firing rates on right trials exceeded those on left trials. The AUROC was calculated as the probability that the firing rate for a unit on a randomly chosen right trial exceeded a randomly chosen left trial, with ties counting as 0.5. AUROC values greater than 0.5 indicated right-selective neurons, while values less than 0.5 indicates left-selective neurons. Statistical significance was assessed using a permutation test (1,000 shuffles), where the trial type label was randomized uniformly for each trial. We defined *P*-values as the proportion of shuffled AUROC values whose absolute deviation from 0.5 was greater than or equal to that of the observed AUROC. Multiple comparisons were corrected for using a Bonferroni adjustment, in which the significance threshold (α=0.05) was divided by the number of epochs tested (n=5 epochs). Selectivity onset was defined for each neuron as the first time point that the AUROC exceeded a directional threshold (≥ 0.65 for right, ≤ 0.35 for left) for at least 50 ms continuously to avoid spurious transients.

### Subspace identification

Movement-null and movement-potent subspaces were identified following the approach described in ref^39^ (**Fig. 2**). Single trial neural activity was first binned in 5 ms intervals and smoothed with a causal Gaussian kernel (20 ms s.d.). Each trial’s activity for each neuron was subtracted by the average baseline activity across trials and scaled by the standard deviation across trials (baseline: –2.1:-1.85 sec before the go cue, during the ITI). **X**_moving_ ∈ ℝ^B_p^ ^×^ ^N^ and **X**_stationary_ ∈ ℝ^B_n^ ^×^ ^N^ were defined using the single trial neural activity when the animal was moving or stationary, respectively. B_n was the number of time bins during which the animal was annotated as stationary and B_p the number of time points annotated as moving. *N* is the number of neurons. **X**_moving_ and **X**_stationary_ contained data from all correct and error trials. Movement-null and movement-potent subspaces were identified through the following optimization problem:

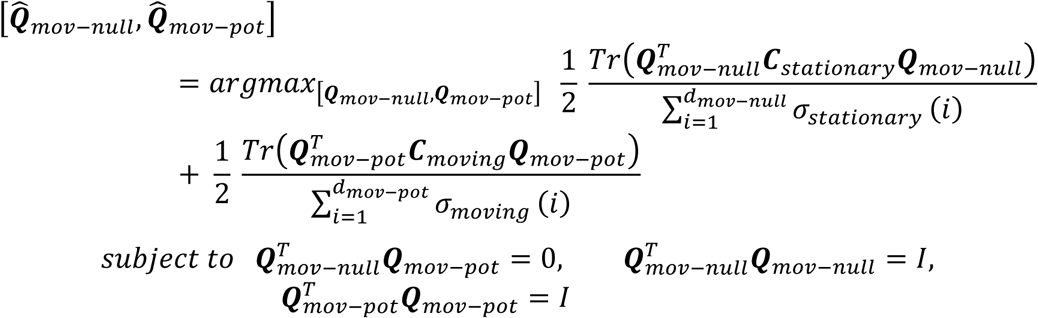

where **C**_stationary_ and **C**_moving_ are the covariances of **X**_stationary_ and **X**_moving_, σ*_stationary_* and σ*_moving_* are the singular values of **C**_stationary_ and **C**_moving_, and d_mov-null_ and d_mov-potent_ are the dimensionality of the subspaces. This optimization problem jointly identifies the subspaces that maximally contain the variance in neural activity during the delay and response epochs. Therefore, the dimensionality of the full population was equal to the dimensionality of the delay or response subspaces for sessions containing forty or fewer neurons. Optimization was performed using the manopt^90^ toolbox for MATLAB.

We quantified the variance explained in a subspace as in refs^19,91^ (**Fig. 2c** and **Fig. 4c**):

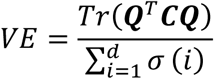

where **Q** is the subspace, **C** is the covariance of neural activity, and σ are the singular values of **C**. This normalization provides the maximum variance that can be captured by *d* dimensions.

Unit activity was reconstructed from movement-null and movement-potent subspaces according to the following equation:

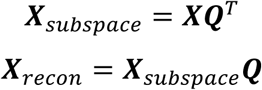

where ***X****_recon_* is the reconstructed neural activity, ***X****_subspace_* is the projected neural activity, ***X*** is single-trial firing rates, and ***Q*** is either the movement-null or movement-potent subspace.

### Dimensionality of subspaces

The dimensionality of movement-null and movement-potent subspaces was determined from the neural activity during stationary and moving time points, respectively. Null distributions were created for each eigenvalue of ***X****_stationary_ and **X**_moving_* by shuffling the time bins separately for each neuron. The remaining covariance structure is then due to chance. This procedure was repeated 1000 times to generate a null distribution for each eigenvalue. The number of eigenvalues of the unshuffled data that exceeded the 95^th^ percentile of the null distribution was used as an upper bound estimate of dimensionality^92^.

### Subspace preference

To quantify the subspace preference index (**Fig. 2e** and **Extended Data** Fig. 2g), we computed a preference index for each neuron based on its activity along experimentally defined subspaces, with values ranging from –1 (movement-potent preference) to +1 (movement-null preference). This was calculated for each neuron as:

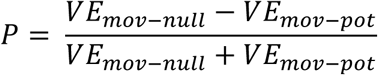

Where *VE* is the variance explained of each individual neuron by the movement-null or movement-potent subspace. *VE* for a neuron was calculated as:

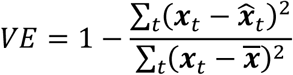

where ***x****_t_* is the single trial firing rate, 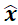 is the firing rate reconstructed from either subspace, and *t* is the time bin.

Subspace preferences towards random dimensions were calculated using a procedure similar to the one described in refs^19,39^. Random subspaces were generated by sampling vectors from a standard normal distribution and aligning them to the covariance structure of the population neural activity using the transformation *VDV^T^ v*, where *V* and *D* are the eigenvectors and eigenvalues of the covariance matrix, and *v* is the random dimension sampled. Sampled dimensions were biased in this way to account for the unequal amounts of variance between moving and stationary time points. The sampled dimensions were normalized and then orthogonalized using MATLAB’s orth() function, which defined the final random subspace. This procedure was repeated 1,000 times, yielding a distribution of preference values for each neuron based on projections into randomly sampled subspaces. Error bars are not shown in **Fig. 2e** because this statistical method generates a distribution for the ‘null hypothesis’ but not error bars.

### UMAP embeddings of neural activity reconstructions

The run_umap() function from the MATLAB File Exchange (https://www.mathworks.com/matlabcentral/fileexchange/71902-uniform-manifold-approximation-and-projection-umap) was used to obtain UMAP embeddings of neural activity reconstructions from either the movement-null or movement-potent subspace (***Fig. 2f,g***, **Fig. 4f *Extended Data Fig. 1***). Train and test sets were obtained by sampling 50% of left and right correct trials each. Before sampling trials to create the train and test sets, left and right trials were size matched by randomly sampling without replacement the minimum of the number of left and right trials. Training and testing data were z-scored per neuron using the mean and standard deviation of activity across all time points and both trial types in the training set. Trial types were concatenated in time, resulting in a matrix of trial-averaged movement-null reconstructions of size (N_time_ * 2, N*_units_*), and provided as training data as input to run_umap() (https://www.mathworks.com/matlabcentral/fileexchange/71902-uniform-manifold-approximation-and-projection-umap) with the following parameters: n_components: 2, n_neighbors: 30, min_dist = 0.3, metric = ‘cosine’. The testing data were then reduced to UMAP space using the learned transformation. For **Fig 2f,g** and **Extended Data** Fig. 1, the training and testing data consisted of both UNs and L-ETNs. For **Fig. 4f**, the training and testing data consisted of both ALM and medulla units.

### Using waveform and autocorrelogram to discriminate L-ETNs and UNs

We used linear discriminant analysis to identify UNs whose extracellular waveform and spiking statistics resemble L-ETNs. For each unit, we randomly sampled 1000 waveforms, or the max number of waveforms, from the extracellular voltage data using a custom MATLAB script. 0.5 ms before and 1 ms after the spike time was extracted. The sampled waveforms were averaged, and peak normalized so that the minimum value was –1. The 3D autocorrelogram (ACG)^93^ was calculated for each neuron using 10 firing rate bins, and firing rates were calculated in 100 ms intervals. Each firing rate bin then consisted of 2D-ACG for a different decile of mean firing rate.

For each neuron, we constructed a feature vector by concatenating principal components of the normalized average waveforms and 3D-ACGs, retaining the top 20 components from each feature, resulting in *N* feature vectors *f* of length 40. *N* is the total number of UNs and L-ETNs. The feature vectors were normalized using the L2 norm. A binary LDA was then trained to discriminate L-ETN feature vectors from UN feature vectors using MATLAB’s fitcdiscr with a linear discriminant. Because only two classes were modeled, the classifier yields a single Fisher axis onto which feature vectors for all units were projected (**Extended Data** Fig. 2e). Any UN whose projection on the Fisher axis exceeded the 5th percentile of the L-ETN projection distribution was labeled “L-ETN-like” and flagged for exclusion. We applied an additional distance criterion: UNs identified from the linear discriminant analysis were only excluded if their estimated location was within 100 µm of channels containing responding L-ETNs, a proxy for being in or near lower layer 5b (see *Determining spatial distance of unidentified neurons to lower layer 5b* below; **Extended Data** Fig. 2a,b).

### Determining spatial distance of unidentified neurons to lower layer 5b

As a proxy for distance of a UN to lower layer 5b, we determined how far in space a UN was from a simultaneously recorded L-ETN. We first determined what channels on our recording electrodes contained L-ETNs. To do this, we quantified the ratio of the amount of variance in a 5 ms window prior to photostimulation onset, and a 10 ms window after photostimulation onset for each channel (**Extended Data** Fig. 2a). We then manually chose a threshold through visual inspection to identify channels which had L-ETNs that reliably responded to photostimulation. Channels were assigned to each UN and L-ETNs based on which channel its waveform was most prominent on (largest amplitude spike). The distance each UN was from the lower layer 5b was defined as the Euclidean distance from its assigned channel to the nearest channel that passed the threshold criteria (**Extended Data** Fig. 2b).

### Neural network models

Neural network models were trained to predict single trial neural firing rates from orofacial movements (**Fig. 3a**). Orofacial movements were provided to the network as input in the form of singular vectors of video data and motion energy data. Video data from both the side view and the bottom view were concatenated so that both movement direction and amplitude could be captured. 100 video and 100 motion energy singular vectors (SV) were obtained through the Facemap GUI^68^. Training data consisted of 70% of trials and the remaining trials were used for testing. Before splitting the data into train/test sets, the number of left and right trials were sampled without replacement equal to the minimum number of trials between the trial types.

The neural network architecture was kept the same as in ref^68^, consisting of an initial one-dimensional convolutional layer that learned ten temporal filters from the input data, followed by a 2-layer feedforward network with ReLU nonlinearity, and a final readout layer that transformed learned features to predictions of single trial firing rates. The feedforward layers had dimensionalities of 50 and 256. The readout layer was a fully-connected layer with output dimensions equal to the number of neurons in a session.

Dropout with a probability of 0.2 was added to the feedforward layers. Individual networks were trained to predict neural activity from each session over 1000 epochs or until the test loss decreased by less than 1 × 10^-5^ for 20 consecutive epochs. The learning rate was initialized at 1 × 10^−2^ and decreased by 10% if the test loss increased for 5 consecutive epochs. Optimization was performed using the AdamW optimizer in PyTorch. Equal numbers of left and right trials were sampled and subsequently divided into an 80/20 train/test split.

The variance explained of a single neuron’s firing rates were calculated as:

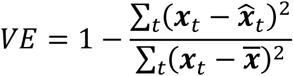

where ***x****_t_* is the single trial firing rate, 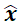 is the predicted firing rate, and *t* is the time bin. The mean squared error between network predictions and single trial firing rates were used to compute the train and test loss at each epoch.

### Evaluating network models against linear models

Network models were evaluated against ridge regression models fit to perform the same task of predicting single trial firing rates from 100 video and 100 ME SVs. Equal numbers of left and right trials were sampled and subsequently divided into a 60/20/20 train/validation/test split. The validation split was used to estimate the optimal ridge parameter. The optimal ridge parameter was then used along with the train set to fit the linear model using MATLAB’s ridge() function. The test split was used to estimate the amount of variance explained of each neuron’s single trial firing as described in the previous section.

### Curation of Mesoscale Activity Map Dataset (MAP)

We used publicly available data (MAP: https://dandiarchive.org/dandiset/000363) consisting of Neuropixels 1.0 and Neuropixels 2.0 recordings across many brain regions simultaneously, including the ALM and medullary reticular nuclei, in mice performing the same delayed-response task as used in this study^58^ (**Fig. 4**). The location in Allen Common Coordinate Framework (CCF) space, along with unit classification based on common spike sorting metrics, were used to identify sessions with at least 30 recorded units per region of interest that were classified as ‘good’. The regions of interest consisted of both hemispheres of the ALM (labeled as MOs in the Allen CCF and described as targeting ‘ALM’ in the data files) or the ALM and the medullary nuclei. Medullary reticular nuclei as defined here consisted of three regions: the intermediate reticular nucleus (IRN), the gigantocellular reticular nucleus (GRN), and the parvicellular reticular nucleus (PARN). The IRN contains circuits responsible for low-level control of orofacial movements^61–63^ and the GRN and PARN border the IRN. We include the GRN and PARN because the probe localization technique used in ref^58^ is precise to approximately 100 µm^94^; units assigned to the GRN and PARN may have been recorded from the IRN. For simultaneous bilateral ALM recordings, we included 20 sessions from 10 mice, consisting of 2016 units (961 from right ALM and 1055 from left ALM). For simultaneous ALM and medulla recordings, we included 11 sessions from 7 mice, consisting of 1328 units (604 from ALM and 724 from medulla).

Spike counts were binned with a 5 ms bin width and smoothed with a causal Gaussian kernel (20 ms s.d) to obtain single-trial firing rates. Movement-null and movement-potent subspaces and projections within subspaces were calculated as described above.

Moving and stationary time points were identified using the provided keypoint tracking data of the jaw, and nose from a camera recording the side profile of the mouse’s face at 300 Hz. The average velocity magnitude of the jaw and nose across time and trials was manually thresholded to classify when a mouse was moving or stationary, per session. If two movement bouts (continuous movements for at least 100 ms) were separated by 50 ms or less, they were merged.

### Cross-region decoding using activity within movement-null and movement-potent subspaces

Cross-region decoding was performed using projections of single-trial firing rates within the movement-null subspace (**Fig. 4d**). For each session, decoding was carried out separately for the two possible source–target configurations (simultaneous ALM and medulla recordings: ALM to medulla and medulla to ALM; cross-hemisphere ALM recordings: right ALM to left ALM and left ALM to right ALM).

At time bin *t*, the goal was to predict the activity of all dimensions in the target region’s subspace from the recent activity of the source region’s subspace. Predictors consisted of the projections from the source region at time bin *t* and the eight preceding bins, corresponding to a 45 ms temporal window (5 ms bins). This produced a predictor matrix of size *N_trials_* × (*D_source_*×9) where *D_source_* is the number of dimensions in the source region’s subspace. The response variable was the matrix of target region activity at time *t* of size *N_trials_* × *D_target_*.

Ridge regression (L2 regularization, *λ* = 0.01) was used to fit the mapping from source to target activity. Model performance was evaluated at each time bin as the variance explained (R^2^) of the target activity on a held-out test set (70% train / 30% test split). Paired *t*-tests were performed on the average variance explained across all task epochs.

### Coding direction analysis

CDs were defined as directions in neural activity state space, defined by firing rates, that maximally separated trajectories of correct lick right and lick left trials.

CD_early_ and CD_late_ were calculated as:

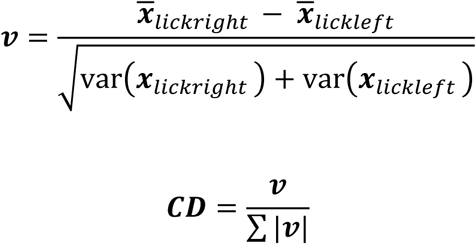

The firing rate for each unit was z-scored using baseline mean and standard deviation (baseline: – 2.15 to –1.85 seconds relative to the go cue, during the inter-trial interval).

For each unit, the mean spike rate difference between right lick trials, 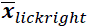, and left lick trials, 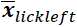, was calculated in either the first 500 ms of the sample epoch (CD_early_) or the last 500 ms of the delay epoch (CD_late_). The vector ***v*** was obtained by first subtracting mean right and left lick firing rates, then normalizing by the square root of the sum of the variances across conditions. Finally, ***v*** was normalized by its L_1_ norm to ensure projections do not scale with the length of ***v***, the number of units simultaneously recorded.

Projections along each CD ∈ ℝ^N^ were obtained by

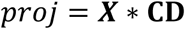

where ***X*** ∈ ℝ^(B*K)^ ^×^ ^N^ is the population activity matrix and B is the number of time bins, K the number of trials, N the number of neurons, and *T* is the transpose operator.

To create matrices of CD correlation at each time bin (**Extended Data** Fig. 3c), CDs were calculated at each time bin from trial-averaged firing rates of all units (5 ms time bins, filtered using a causal Gaussian kernel with 20 ms s.d.). CDs were normalized by their Euclidean norms to produce unit vectors. Correlation matrices were then obtained as the inner product of CD vectors at each pair of time points.

### Hierarchical bootstrapping

Projections along CDs were obtained via a hierarchical bootstrapping procedure^95,96^ (**Extended Data** Fig. 3). Pseudopopulations were constructed by randomly sampling with replacement *M* mice, *S*_M_ sessions per sampled mouse, 50 correct trials of each type, and *N*_S_ neurons. *M* is the number of mice in the original cohort*, S*_M_ the number of sessions per mouse, and *N*_S_ the number of neurons in each session. Bootstrapping was repeated for 1,000 iterations. *M*ean and 5–95% confidence intervals (*shaded regions*) of selectivity along projections are shown to indicate where projections significantly differ from zero (*p* < 0.05, one-sided test, bootstrap; ***Extended Data Fig. 3a,b***).

### Statistics and reproducibility

No statistical methods were used to predetermine sample sizes. All *t*-tests were two-sided unless stated otherwise. All animals from which electrophysiology and behavior data were collected were included in the study. Trial type was randomly determined by a computer program during electrophysiology sessions. Experimenters were blind to trial type during all data collection and spike sorting.

## Data availability

Data will be made publicly available upon publication.

## Code availability

All custom code used for analyses will be made available on Github upon publication. Code for identifying movement-null and movement-potent subspaces is available at https://github.com/economolab/subspaceID.

## Acknowledgements

We thank J. Birnbaum and O. Elsayed for comments on the manuscript. This work was supported by the Whitehall Foundation, the Klingenstein Fund, the Simons Foundation and National Institutes of Health R01NS121409 and U19NS137920.

## Contributions

M.A.H and M.N.E conceived the project. M.A.H, Y.H, and M.N.E designed the experiments. M.A.H, and Y.H performed experiments. M.A.H and M.N.E analyzed data. M.A.H. and M.N.E wrote the manuscript. M.N.E supervised research.

